# Balancing Speed and Precision in Protein Folding: A Comparison of AlphaFold2, ESMFold, and OmegaFold

**DOI:** 10.1101/2025.06.20.660709

**Authors:** Anna Hýskova, Eva Maršálková, Petr Šimeček

## Abstract

We compared the performance of three widely used protein structure prediction tools—AlphaFold2, ESMFold, and OmegaFold—using a dataset of over 1,300 newly created records from the PDB database. These structures, resolved between July 2022 and July 2024, ensure unbiased evaluation, as they were unavailable during the training of these tools. Using metrics such as root mean square deviation (RMSD), template modeling score (TM-score), and predicted local distance difference test (pLDDT), we found that AlphaFold2 consistently achieves the highest accuracy but depends on high-quality sequence alignments. In contrast, ESMFold and OmegaFold provide faster predictions and excel in challenging cases, such as rapidly evolving or designed proteins with limited sequence homology. Comparing ESMFold and OmegaFold, ESMFold achieves higher confidence scores (pLDDT) and structural similarity (TM-score). OmegaFold is competitive in specific contexts, such as de novo-designed proteins or sequences with limited evolutionary information. Additionally, we demonstrate that machine learning models trained on protein language model embeddings and pLDDT confidence scores can predict potential structure prediction failures, helping to identify challenging cases early in the pipeline.

## Introduction

All living organisms—from simple bacteria and algae to plants, fungi, animals, and humans—contain a multitude of proteins that participate in virtually every cellular process [1, 2]. These molecular machines must fold into specific three-dimensional structures, organized hierarchically at four distinct levels (Fig. 1): from the linear sequence of amino acids (primary structure), through local folding patterns of *α*-helices and *β*-sheets (secondary structure), to the complete three-dimensional arrangement of these elements (tertiary structure), and finally to the assembly of multiple chains into functional complexes (quaternary structure). While the amino acid sequence alone determines the final structure, protein misfolding often leads to disease [3]. Experimental structure determination through X-ray crystallography, cryo-EM, or NMR spectroscopy remains the gold standard [4, 5, 6], but these methods are time-consuming, expensive, and not always feasible. This creates an urgent need for reliable computational prediction methods, particularly as the gap between known protein sequences and solved structures continues to widen—with over 254 million sequences known (UniProtKB) but only about 230,444 experimentally determined structures available in the Protein Data Bank (as in January, 2025).

**Fig. 1.**
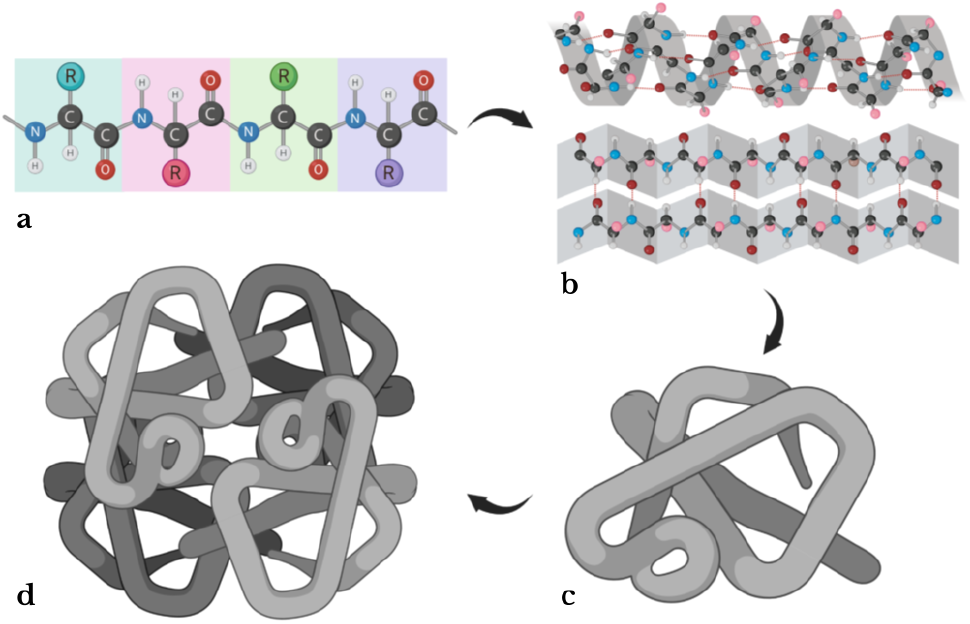
Hierarchical organization of protein structure. (a) Primary structure: the linear sequence of amino acids. (b) Secondary structure: local conformations including *α*-helices and *β*-sheets stabilized by hydrogen bonds. (c) Tertiary structure: the complete three-dimensional fold. (d) Quaternary structure: assembly of multiple chains into functional complexes.

The field of protein structure prediction has been transformed by artificial intelligence approaches. The introduction of AlphaFold2 in 2020 marked a watershed moment, achieving near-experimental accuracy [7]. This success has spurred the development of alternative approaches, particularly language model-based predictors like ESMFold and OmegaFold that can generate predictions without requiring multiple sequence alignments [8, 9]. These newer methods promise faster predictions and potentially better performance on challenging targets like designed or rapidly evolving proteins.

Despite these advances, the field lacks a comprehensive comparison of these tools’ performance on truly novel proteins—structures solved after the tools’ training cutoff dates [10]. Such evaluation is crucial for understanding each method’s strengths and limitations, particularly as these tools become increasingly integrated into structural biology workflows. While the Critical Assessment of Structure Prediction (CASP) [11] and Continuous Automated Model EvaluatiOn (CAMEO) [12] provide valuable benchmarks, they are limited to participating methods and may not reflect real-world usage patterns.

Here, we present a systematic comparison of AlphaFold2, ESMFold, and OmegaFold using a dataset of over 1,300 protein structures deposited in the PDB between 2022 and 2024. Using multiple evaluation metrics including RMSD [13], TM-score [14], and pLDDT [15], we assess both overall performance and specific challenging cases. Our analysis reveals that while AlphaFold2 achieves the highest average accuracy, ESMFold and OmegaFold excel in particular niches, especially for proteins with limited homology information. We also identify protein families and structural features that correlate with prediction success or failure, providing practical guidance for the structural biology community.

## Methods

### Dataset

We compiled a benchmark dataset of 1,327 protein structures deposited in the Protein Data Bank (PDB) between July 2022 and July 2024. This temporal restriction ensures no overlap with training data used by AlphaFold2 (cutoff April 2020), ESMFold (June 2020), or OmegaFold (2021). The dataset contains three distinct groups: (1) single-chain monomers (980 structures), (2) small multi-chain complexes (245 structures with 2-6 chains), and (3) de novo designed proteins whose sequence does not naturally occur in any living organism (102 structures).

Structures were selected using the RCSB PDB Search API [16, 17] with the following criteria: (i) deposition date between July 2022 and July 2024, (ii) protein-only structures without nucleic acids or oligosaccharides, (iii) chain lengths between 20 and 400 amino acids to ensure compatibility with all prediction tools, and (iv) availability of structural information in PDB format. To ensure diversity, structures within monomer and de novo protein groups were filtered to have at most 70% pairwise sequence identity.

We developed a custom PDB file parsing pipeline to extract complete amino acid sequences and experimental C_*α*_ coordinates. The pipeline addresses common challenges in PDB files, including non-standard residue numbering, insertion codes, and post-translational modifications. For modified residues, we reconstructed the original amino acid sequence using BioPython’s extended residue dictionary and MODRES records. Structures containing non-standard residues without clear mapping to canonical amino acids (26 cases) were excluded from the analysis.

Each structure was annotated with protein family classifications using UniProt and PDBe APIs to map PDB identifiers to Pfam and InterPro database entries. These annotations enable analysis of prediction tools’ performance across different protein families and structural motifs. The numbers of protein structures the dataset contained in various stages of the experiment are stated in Table 1. The final curated dataset, including all protein sequences, structures, and family annotations, is available at Hugging Face Hub repository.

**Table 1.** Size of the dataset in various stages of the experiment.

| Group | PDB hits | Selected structures | Successfully processed | Evaluated chains |
| --- | --- | --- | --- | --- |
| Monomers | 3830 | 1000 | 980 | 979 |
| Small Complexes | 3988 | 250 | 245 | 255 |
| De Novo Proteins | 139 | 103 | 102 | 102 |
| In Total | 7957 | 1353 | 1327 | 1336 |

(A) Number of hits in PDB database. (B) Number of randomly selected structures. (C) Number of structures with PDB files successfully preprocessed. (D) Final number of chains used in the evaluation dataset (successfully predicted by all three tools).

### Structure Prediction Tools

Three tools were selected for protein structure prediction: AlphaFold2, ESMFold, and OmegaFold. While alignment-based AlphaFold2 is an obvious choice, considering how widely used it is [10], language model-based ESMFold and OmegaFold were chosen because they provide promising results with much lower requirements on time and computational power, making them more suitable for large-scale applications [8, 9].

#### AlphaFold2

We used AlphaFold v2.1.1 running on the university e-INFRA CZ infrastructure with its monomer model and reduced database settings to optimize computational resources. The model architecture consists of two main components: (i) an Evoformer module, which processes multiple sequence alignments (MSAs) and pairwise representations through 48 transformer blocks, and (ii) a structure module that converts the refined representations into 3D coordinates through 8 equivariant transformer blocks with Invariant Point Attention. MSAs were generated using Uniref90, BFD, and MGnify databases. For each sequence, five model predictions were generated and ranked by predicted confidence, with the highest-confidence model (ranked 0.pdb) selected for evaluation.

#### ESMFold

Predictions were obtained via REST API calls to the ESM Metagenomic Atlas. ESMFold combines two components: (i) the ESM-2 protein language model with 15B parameters, pre-trained on masked sequence prediction, and (ii) a folding head consisting of 48 folding blocks that process sequence and pairwise representations. Unlike AlphaFold2, ESMFold predicts structures directly from single sequences without requiring MSA generation.

#### OmegaFold

Predictions were performed using OmegaFold v1.0 running on university computational cluster with NVIDIA A40 GPU. OmegaFold employs: (i) OmegaPLM, a 670M parameter language model trained on masked protein sequences, and (ii) a Geoformer architecture that refines the language model representations to be geometrically consistent before structure prediction. Like ESMFold, OmegaFold operates on single sequences without MSA requirements.

All predictions were made for individual protein chains, as both ESMFold and OmegaFold do not support prediction of protein complexes. While AlphaFold2 offers a multimer model, we used its monomer model to ensure fair comparison. Source code, configuration files, and prediction outputs are available at GitHub repository.

### Evaluation Metrics

We employed three complementary metrics to assess prediction quality: RMSD measuring atomic distance deviation, TM-score evaluating topological similarity, and pLDDT reflecting model confidence.

#### Root Mean Square Deviation (RMSD)

RMSD quantifies the average distance between corresponding C_*α*_ atoms in superimposed structures:

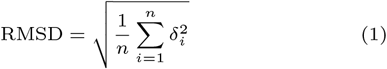

where *n* is the number of aligned C_*α*_ atom pairs and *δ*_*i*_ is the distance between atoms in the *i*-th pair. To compute RMSD, we first extract C_*α*_ coordinates from both experimental and predicted structures, then determine the optimal superposition using the Bio.SVDSuperimposer module from BioPython [18], which finds the rotation and translation matrices minimizing the RMSD value. While RMSD is widely used, it is sensitive to protein size and can be disproportionately affected by local structural deviations.

#### Template Modeling Score (TM-score)

TM-score evaluates the topological similarity of protein structures while accounting for protein length:

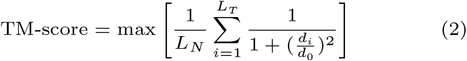

where *L*_*N*_ is the length of the reference structure, *L*_*T*_ is the number of aligned residues, *d*_*i*_ is the distance between the *i*-th pair of aligned residues after superposition, and 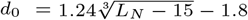 is a length-dependent scaling factor. TM-score ranges from 0 to 1, with values above 0.5 indicating proteins share the same fold and 1 representing perfect structural alignment. Unlike RMSD, TM-score is length-normalized and less sensitive to local structural variations.

#### Predicted LDDT (pLDDT)

The predicted local distance difference test (pLDDT) is a confidence metric provided by each prediction tool. For each residue, it estimates the expected agreement between predicted and experimental structures on 0 to 100 scale. Scores above 90 indicate high prediction confidence. Scores above 70 suggest at least reliable backbone prediction.

For our analysis, we used the mean pLDDT across all residues in each protein chain. While pLDDT correlates with prediction accuracy, high confidence scores do not guarantee correct structure prediction, particularly for challenging targets like intrinsically disordered regions or proteins with limited homology information.

### Statistical Analysis and Annotation

We compared these metrics across our dataset using Kruskal-Wallis tests followed by Dunn’s method with Bonferroni correction for multiple comparisons. The correlation between metrics was assessed using Spearman’s rank correlation coefficient.

Protein chains were mapped to functional annotations using UniProt and PDBe APIs. For family-specific analysis, we focused on Pfam and InterPro families with at least 10 member proteins in our dataset. The experimental method of structure determination (X-ray crystallography, cryo-EM, or NMR) was recorded for each chain to assess potential biases in prediction accuracy.

Predictions were classified as “poor” if they met any of the following criteria: average pLDDT *<* 70, TM-score *<* 0.5, or RMSD *>* 9 Å. The 9 Å RMSD threshold was chosen to match the resolution cutoff used in training AlphaFold2. Statistical significance of family-specific enrichment in poor predictions was assessed using Fisher’s exact test with Benjamini-Hochberg correction for multiple comparisons.

### Implementation and Availability

All analysis code was implemented in Python using BioPython for structure manipulation, tmtools for TM-score calculation, and scipy.stats for statistical testing. The complete dataset, including protein sequences, experimental structures, predictions, and evaluation results is available at https://huggingface.co/datasets/hyskova-anna/proteins. Source code and documentation are provided at https://github.com/ML-Bioinfo-CEITEC/CAoPSPT.

## Results

The structure of 1,337 protein chains was predicted using AlphaFold2, ESMFold, and OmegaFold. Of these, AlphaFold2 failed to predict a single chain (8B2M:A), which was excluded from the evaluation; all remaining predictions were obtained successfully. Selected examples of predictions aligned with their experimental structures are visualized in Figure 2.

**Fig. 2.**
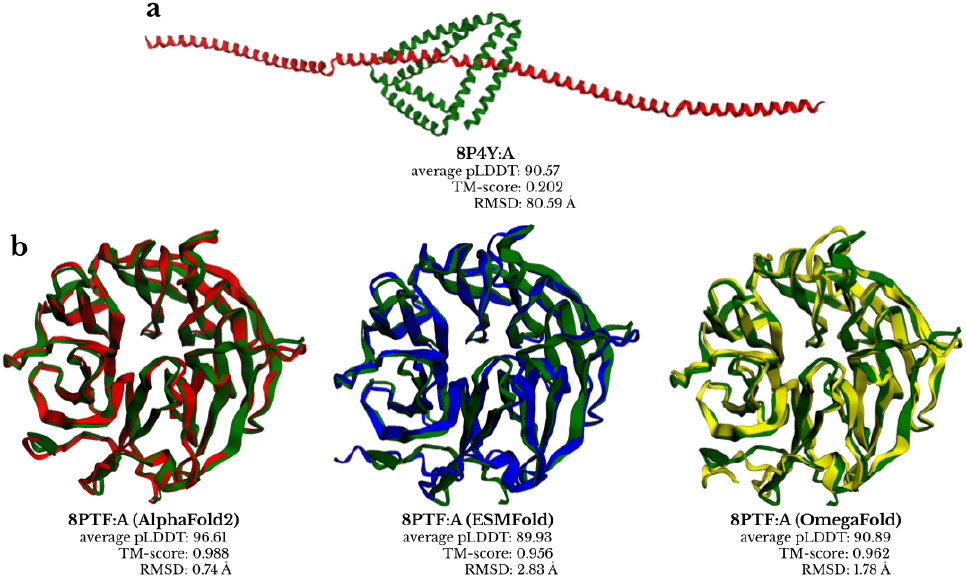
Examples of structure predictions from AlphaFold2 (red), ESMFold (blue) and OmegaFold (yellow) aligned with corresponding experimentally determined structures (green). (a) An example of a poorly predicted structure (8P4Y:A) by AlphaFold2. (b) Structure of protein 8PTF:A showing varying prediction quality across tools.

### Comparative Performance Analysis

All three tools demonstrated generally satisfactory performance, with AlphaFold2 achieving the highest accuracy across all metrics (Figure 3). AlphaFold2 predictions showed the highest median TM-score (0.96) and lowest median RMSD (1.30Å), followed by ESMFold (TM-score: 0.95, RMSD: 1.74Å) and OmegaFold (TM-score: 0.93, RMSD: 1.98Å). Consistently, AlphaFold2 displayed the highest confidence in its predictions with median pLDDT of 92.65, compared to 87.40 for ESMFold and 89.00 for OmegaFold.

**Fig. 3.**
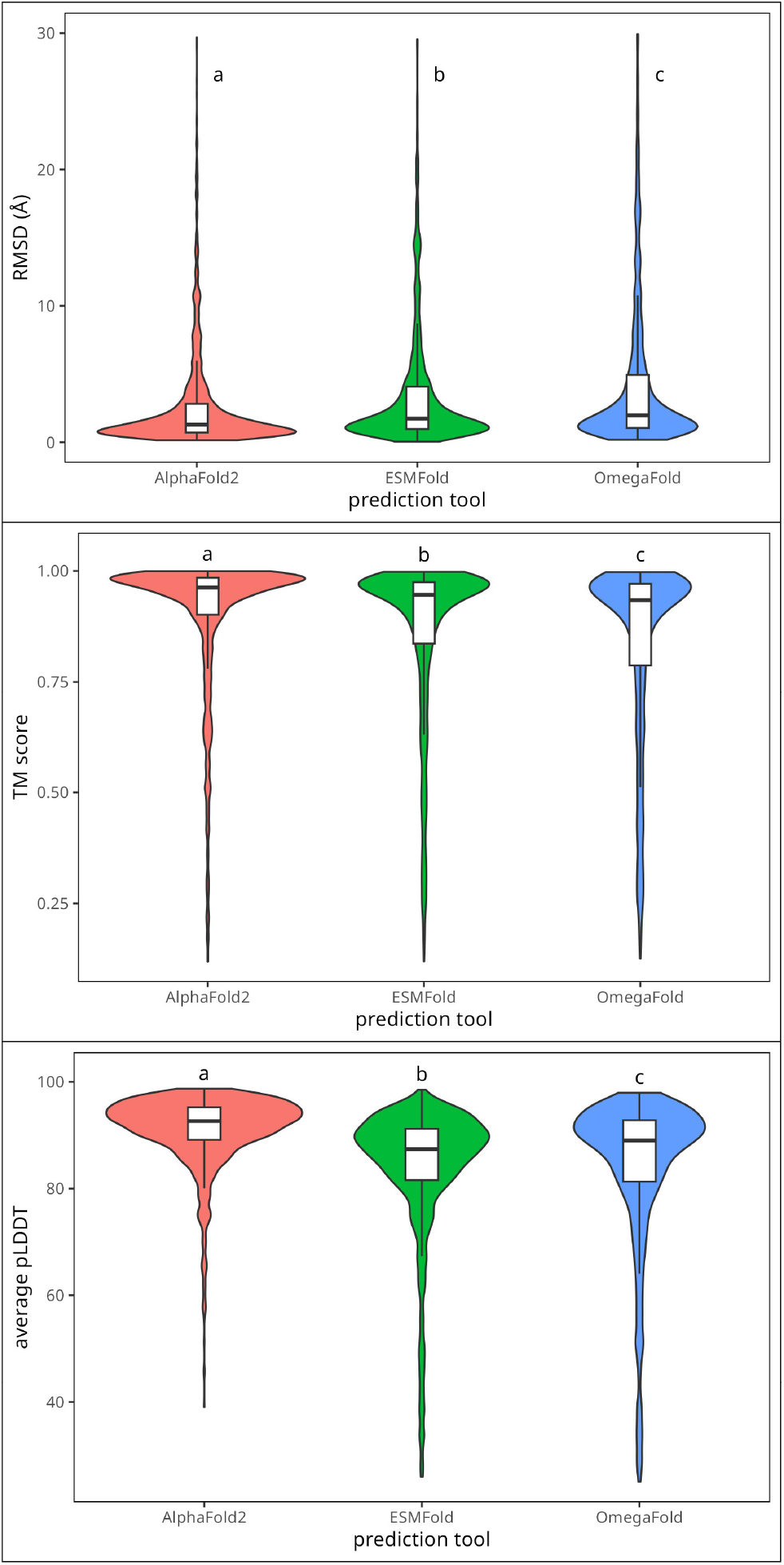
Performance comparison across prediction tools. Distribution of (a) RMSD values, (b) TM-scores, and (c) pLDDT scores. Box plots show median, quartiles, and outliers. All pair comparisons have been statistically significant (*p <* 0.01).

### Metric Correlations and Their Dependencies on Sequence Length And Other Factors

We observed significant correlations between prediction confidence (pLDDT) and accuracy metrics (Figure 4). Most notably, there was a negative correlation between average pLDDT and RMSD (Spearman’s *ρ* = *−*0.87, *−*0.87, and *−*0.88 for AlphaFold2, ESMFold, and OmegaFold, respectively) and a positive correlation between average pLDDT and TM-score (*ρ* = 0.60, 0.66, and 0.71). The correlation was strongest for ESMFold and OmegaFold, suggesting that their confidence scores more accurately reflect prediction quality than those of AlphaFold2.

**Fig. 4.**
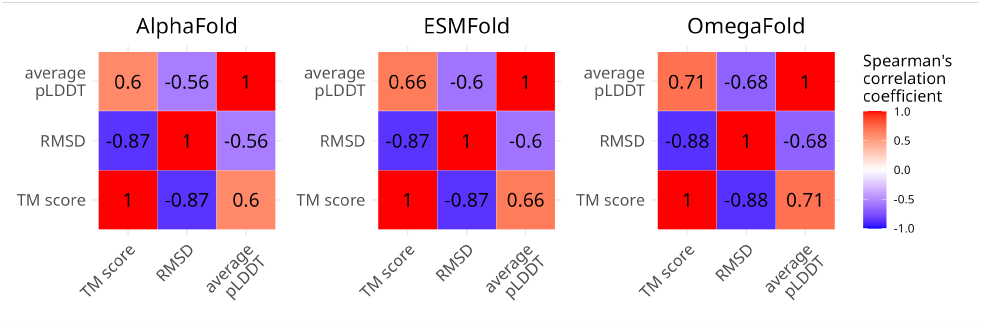
Correlation analysis between prediction metrics. Heatmaps show Spearman’s correlation coefficients between average pLDDT, RMSD, and TM-score for each prediction tool. All correlations are statistically significant (*p <* 0.001).

While low-confidence predictions rarely achieved good accuracy metrics, we found numerous cases of incorrect structures with high pLDDT scores across all tools (Figure 5)

**Fig. 5.**
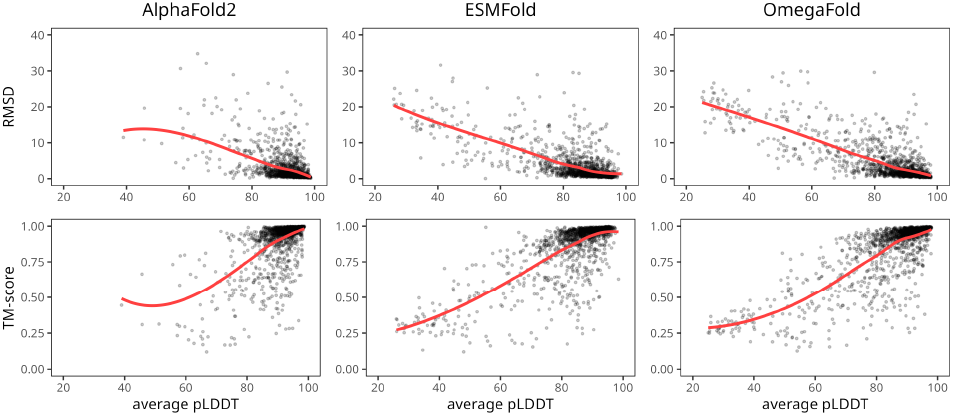
Dependency of RMSD and TM-score on average pLDDT of structures generated by different tools. The LOESS curve (red) was obtained by locally estimated scatterplot smoothing. Sample points with RMSD greater than 40 Å are omitted from the visualization for better clarity.

Analysis of sequence length dependency also revealed interesting patterns. While RMSD showed weak correlation with sequence length, TM-scores displayed stronger positive associations, particularly for AlphaFold2 (*ρ* = 0.41, *p <* 0.001). This suggests that predictions for shorter proteins (*<* 100 amino acids) tend to achieve lower TM-scores across all tools, though this trend is less pronounced in RMSD values due to the metric’s inherent length dependency. ESMFold and OmegaFold showed weaker but still significant correlations with sequence length (*ρ* = 0.29 and *ρ* = 0.28, respectively, for TM-score).

The experimental method used for structure determination significantly influenced prediction accuracy (Figure 7). All tools performed best on X-ray crystallography structures (median RMSD: 1.24Å, 1.65Å, and 1.89Å for AlphaFold2, ESMFold, and OmegaFold, respectively) but struggled with NMR-determined structures (median RMSD: 2.31Å, 2.89Å, and 3.12Å). This pattern likely reflects both the inherent flexibility of proteins amenable to NMR analysis and the predominance of X-ray structures in training data.

**Fig. 6.**
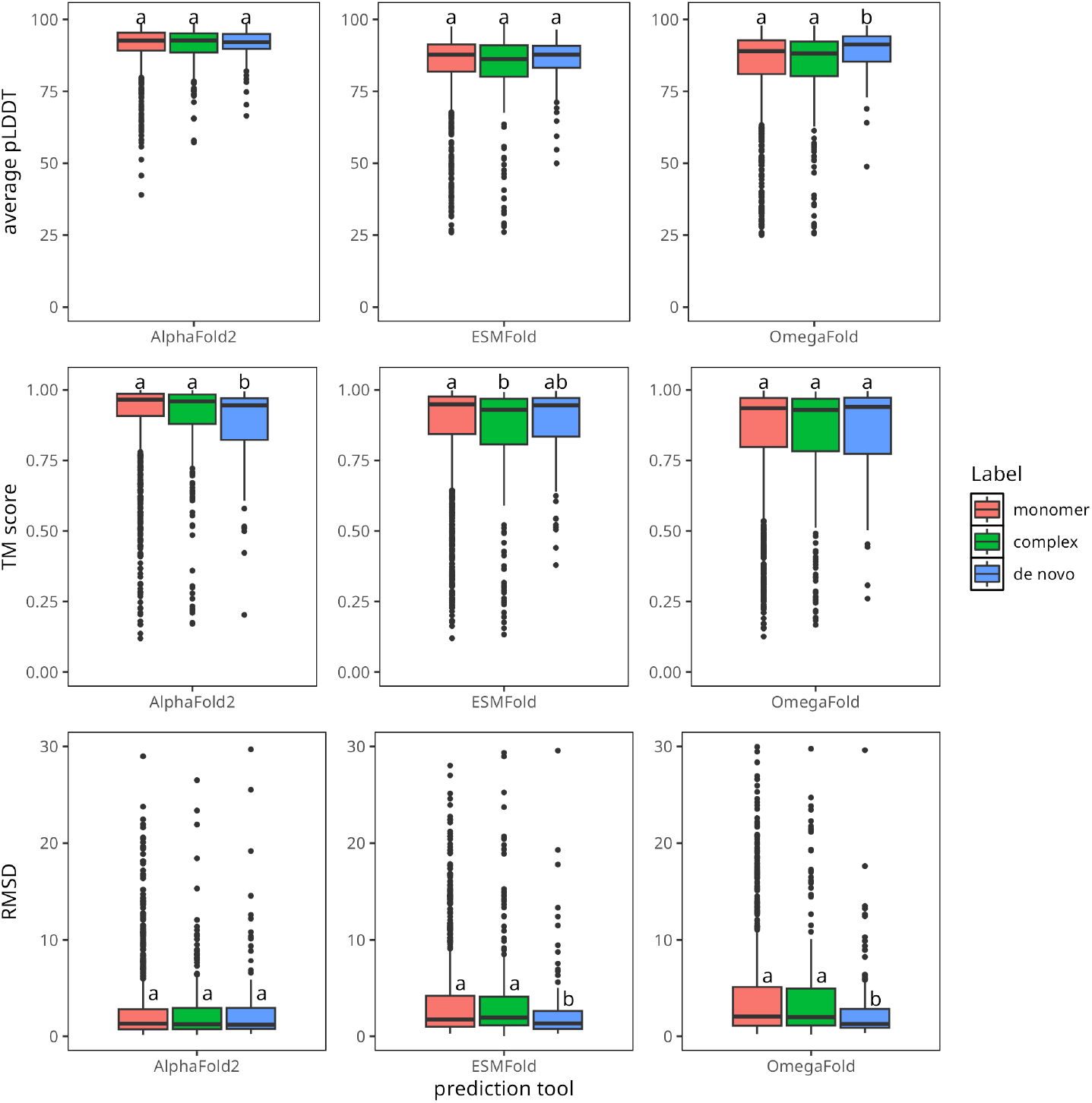
Dependency of average pLDDT, TM-score, and RMSD on the type of protein chain being predicted. The differences between groups were tested by Kruskal-Wallis test, post-hoc comparisons were done using Dunn’s method with a Bonferroni correction for multiple tests. Statistical significance visualized by difference in letter codes. Sample points with RMSD greater than 30 Å are omitted from the visualization for better clarity.

**Fig. 7.**
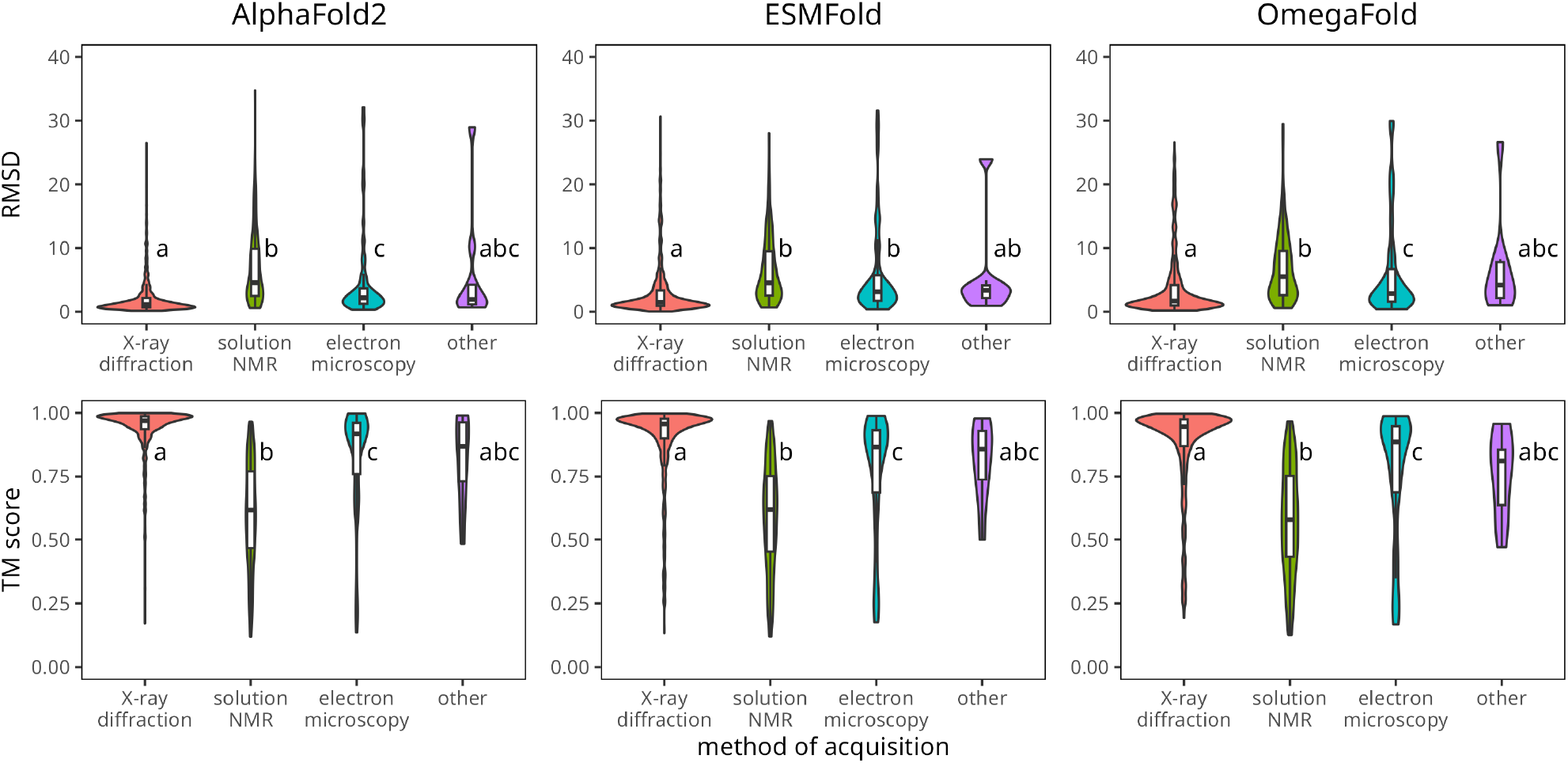
Dependency of RMSD and TM-score on the experimental method of acquisition of the protein chain structure. The differences between groups were tested by Kruskal-Wallis test, post-hoc comparisons were done using Dunn’s method with a Bonferroni correction for multiple tests. Statistical significance visualized by difference in letter codes. Sample points with RMSD greater than 40Å are omitted from the visualization for better clarity.

When comparing performance across different protein types (monomers, complexes, and de novo proteins), we observed an interesting pattern, see Figure 6. While all tools generally performed similarly across these categories, there are two notable exceptions. First, ESMFold and OmegaFold achieved significantly lower RMSD values for de novo proteins compared to natural proteins. Second, AlphaFold2 showed a unique weakness with de novo proteins, achieving significantly lower TM-scores for these proteins compared to monomers and complexes. This suggests that language model-based tools may have an advantage in predicting structures of artificial proteins where evolutionary information is limited.

### Analysis of Prediction Failures

We classified predictions as incorrect if they met any of the following criteria: average pLDDT *<* 70, TM-score *<* 0.5, or *RMSD >* 9Å. AlphaFold2 produced the fewest incorrect predictions (8.9% of total), followed by ESMFold (13.0%) and OmegaFold (16.8%). The overlap of prediction failures between tools was limited, suggesting complementary strengths (Figure 8).

**Fig. 8.**
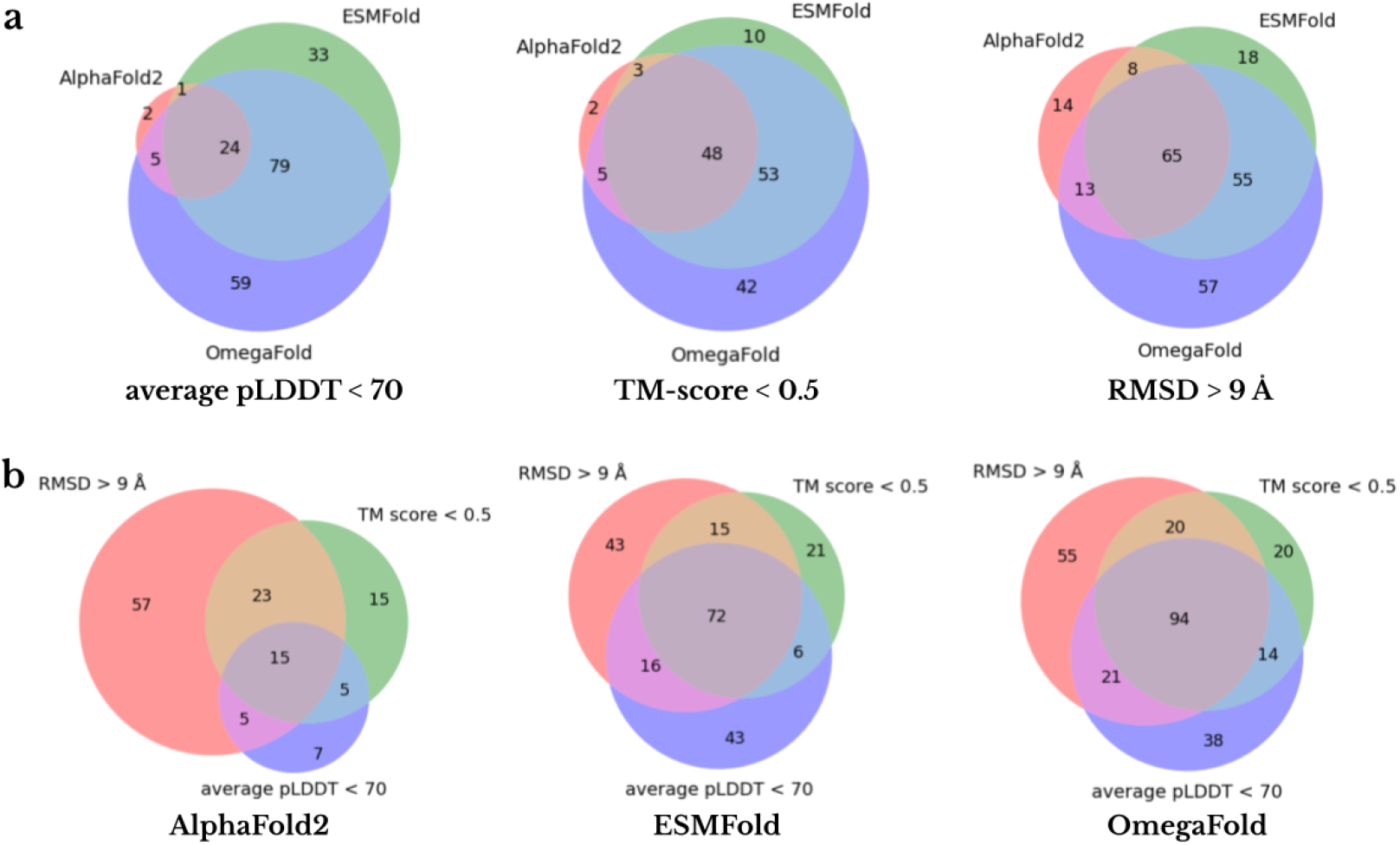
Venn diagrams comparing the overlap of poorly predicted protein chains based on three evaluation criteria: (a) average pLDDT *<* 70, TM-score *<* 0.5, and RMSD *>* 9 Å for AlphaFold2, ESMFold, and OmegaFold, and (b) the overlap of predictions that fail across the three metrics for each tool individually.

Analysis of protein families revealed that proteins lacking Pfam annotations were particularly challenging for AlphaFold2 but not for ESMFold or OmegaFold, highlighting the importance of evolutionary information in AlphaFold2’s predictions. Conversely, viral proteins, especially from coronavirus, were better predicted by AlphaFold2 than by the language model-based tools. All tools showed reduced accuracy for proteins containing leucine-rich repeats or von Willebrand factor A-like domains, suggesting these structural motifs pose particular challenges for current prediction methods.

The analysis of protein family associations revealed distinctive patterns in prediction accuracy. Notably, AlphaFold2 showed significantly reduced performance for proteins lacking Pfam family annotations (odds ratio = 0.67, *p <* 0.01), while ESMFold and OmegaFold maintained consistent performance regardless of family assignments. This pattern was also observed with InterPro annotations, highlighting AlphaFold2’s dependence on evolutionary information.

Certain protein families were consistently well-predicted across all tools. These included protein kinase domains (PF00069, IPR000719), the SH2 domain (IPR000980), and the NAD(P)-binding domain superfamily (IPR036291). Conversely, all tools struggled with leucine-rich repeats (IPR001611, IPR003591) and von Willebrand factor A-like domains (IPR036465), suggesting these structural motifs remain challenging for current prediction methods.

Interestingly, several protein families showed tool-specific prediction patterns. AlphaFold2 excelled at predicting viral protein families, particularly the viral RNA-dependent RNA polymerase (PF00680, IPR001205) and coronavirus-specific proteins (PF05409, IPR043503), achieving significantly better accuracy than ESMFold or OmegaFold (*p <* 0.001). Conversely, the S-adenosyl-L-methionine-dependent methyltransferase superfamily (IPR029063) showed markedly different prediction quality between AlphaFold2 (odds ratio = 1.83, *p <* 0.05) and the language model-based tools (odds ratio = 0.64 and 0.51 for ESMFold and OmegaFold respectively, *p <* 0.001).

### Prediction of Structure Determination Success Using Machine Learning

To help identify potential failures in structure prediction, we trained gradient boosting LightGBM [19] models for AlphaFold2, ESMFold, and OmegaFold, respectively, using ProtBert BFD embeddings [20] calculated from protein sequences and pLDDT scores. The models were trained to predict whether a structure prediction would likely be unsuccessful, allowing early identification of challenging cases. The trained models and source code are available on GitHub repository, enabling to assess potential challenges early in structure prediction pipelines.

### Prediction of Structure Determination Success Using Machine Learning

To help identify potential failures in structure prediction, we trained gradient boosting LightGBM [19] models for AlphaFold2, ESMFold, and OmegaFold, respectively, using ProtBert BFD embeddings [20] calculated from protein sequences and pLDDT scores. The models were trained to predict whether a structure prediction would likely be unsuccessful, allowing early identification of challenging cases. The trained models and source code are available on GitHub repository, enabling to assess potential challenges early in structure prediction pipelines.

As shown in Figure 9, it is typically pLDDT, the length of the sequence, and a few selected embedding elements that have the greatest influence on prediction.

**Fig. 9.**
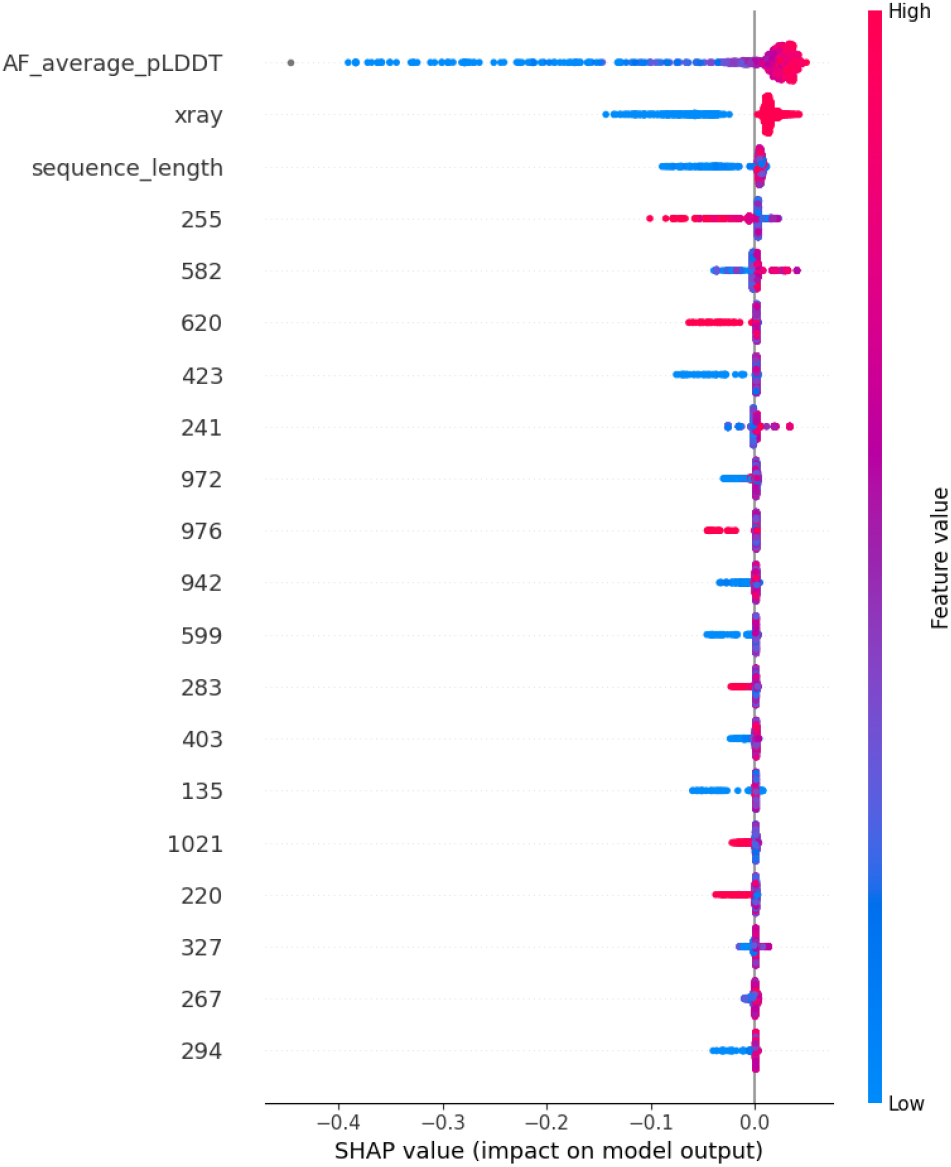
SHAP values of LightGBM model for AlphaFold2.

## Discussion

Since the beginning of this decade, structural biology and protein structure prediction fields have undergone a significant transition. Currently, there are two large projects dealing with this issue: CASP [11] and CAMEO [12]. While AlphaFold2 has participated in both CASP14 and CAMEO, ESMFold has entered only CASP15, and OmegaFold has not been included in either. There are also a few publications dealing with the comparison of protein structure prediction tools, but they usually focus mainly on AlphaFold2 and similar tools (e.g. ColabFold) [21] or perform the evaluation on a particular set of proteins, namely human proteins [22], snake venom toxins [21], and nanobodies [23]. This paper tries to increase our understanding by creating an inclusive dataset of protein structures recently added to PDB.

The key finding of this work is that AlphaFold2 outperforms ESMFold and OmegaFold on a majority of proteins in the dataset, measured by both RMSD and TM-score. When comparing the two protein language-based models, ESMFold seems to be a slightly better choice, as it produced fewer incorrect structures than OmegaFold and achieved significantly better median RMSD and TM-score. Still, the difference in performance between ESMFold and OmegaFold is much smaller compared to the gap between both of these tools and AlphaFold2.

While all three tools rarely produce a good prediction with low confidence, wrong structures with a high average pLDDT are outputted quite frequently. Our analysis revealed that prediction accuracy is influenced by various factors. All three tools performed best when predicting proteins whose experimental structure was determined by X-ray crystallography, while structures determined by NMR proved to be the most challenging. Because NMR is typically used to determine the structures of small proteins, a corresponding decrease in prediction accuracy is observed for shorter sequences.

Interestingly, proteins without family annotations proved particularly difficult for AlphaFold2 but did not change the performance of ESMFold and OmegaFold. A possible explanation is that proteins belonging to no family lack homologs with a known structure, which AlphaFold2 could use as a template during the prediction. In contrast, ESMFold and OmegaFold do not rely on MSAs and modelling templates, so their performance remained largely unaffected.

Our analysis shows several key insights, yet certain constraints of our study must be noted. First, the dataset does not contain only proteins whose experimental structure was previously unknown but also proteins that were just recently analyzed again, usually in different conditions. This might be an advantage for AlphaFold2, which uses a reduced PDB database for template searching during the prediction process. Moreover, the whole analysis focuses only on single protein chains without the context of their interacting partners, which might be crucial for structure formation, especially in protein complexes. Last but not least, all the protein chains in the dataset have a maximum length of 400 amino acids due to using ESMAtlas API.

The field continues to evolve rapidly. Recently, DeepMind released AlphaFold3 [24], followed by several replication efforts including Boltz-1 [25] and HelixFold3 [26]. The code has been made available for noncommercial research purposes in November 2024 [27]. In the future, it would be interesting to predict the structure of whole protein complexes using these newer models and compare the results with predictions of single chains. The analysis could also be extended by including protein chains of all lengths, exploring additional metrics such as the variance of pLDDT, or testing newer models as they become available.

## Author contributions statement

A.H. was responsible for dataset preparation and the majority of coding. A.H. and P.S. collaborated on writing the manuscript. E.M. and P.S. provided critical manuscript review and oversaw project management.

## Acknowledgments

The project was supported by the OPUS LAP program of the Czech Science Foundation, project no. 23-04260L “Biological code of knots – identification of knotted patterns in biomolecules via AI approach”). Computational resources were supplied by the project “e-Infrastruktura CZ” (e-INFRA CZ LM2018140) supported by the Ministry of Education, Youth and Sports of the Czech Republic.

